# LEAPING A CONSERVATION GENETICS GAP: COMBINING POPULATION GENETICS AND SOCIAL NETWORK ANALYSIS FOR *LUPINUS PERENNIS* RESTORATION

**DOI:** 10.64898/2026.08.31.748316

**Authors:** Cooper Kimball-Rhines, Scarlet Taveras-Guzman, Georgia Mavrommati, Brook T. Moyers

**Author notes:** **Corresponding Author**: Cooper Kimball-Rhines.

## Abstract

Genomic data can assist conservation planning, but implementation of genetic-based recommendations is rare. Closing this “conservation genetics gap” is crucial for restoration and translocation decisions, particularly for highly fragmented ecosystems like pine barrens. We first analyze the conservation genetics gap in pine barren management organizations using a survey. We then analyze whole-genome data to characterize population structure and diversity in the pine barren keystone plant *Lupinus perennis*. We find little genetic difference among *L. perennis* populations in the Northeast United States. While populations show signs of isolation by distance (*⍴* = 0.785, *p* = 0.001), we find no correlation between population size and heterozygosity (Estimate = 0.003, *p* = 0.357) or inbreeding (Estimate = 0.028, *p* = 0.416). We also find no effect of seed source population on forecasted effective population size in restoration simulations. These findings and our recommendation that managers consider non-local seed provenancing for *L. perennis* restorations in the Northeast have been integrated into SWAP policy in two states. We generally recommend practitioners record seed provenance, save genetic material for future testing, and incorporate fitness metrics into restoration monitoring. To effectively close the conservation genetics gap, researchers providing conservation recommendations must consult with practitioners before designing experiments.

## Introduction

Inland pine barrens are a globally rare ecosystem that have experienced extreme habitat loss and fragmentation over the last centuries. The natural history and ecology of pine barrens, which are characterized by harsh soils and frequent fires that reset the system’s successional cycle, made this ecosystem a common target of urbanization (Goldstein 1978). Areas of once sprawling pine barren habitat were selected as sites of urban centers by European colonizers, including what are now known as New York City, Albany, NY, Concord, NH, and Washington, DC (Lee 2019). Continued human development and fire suppression policies have led to the loss of >90% of historic pine barrens (Pavlovic 2009). Current pine barren conservation efforts focus on restoring historic fire regimes, often planting or supplementing fire-adapted species after prescribed burns (Clark 2015; Gobster 2021; Petitta et al. 2024).

One such fire-adapted species is *Lupinus perennis* (Fabacaea), commonly known as sundial lupine, wild blue lupine, and perennial lupine (Haines et al. 2011). *Lupinus perennis* is well adapted to recolonizing freshly burned habitat either vegetatively through its robust taproot system or using its heat-triggered cannon-like seed dispersal mechanism (Plenzler 2015). The species was once common throughout its range spanning the East Coast to the Midwest of North America but is now considered at-risk, threatened, endangered, or extirpated in fourteen U.S. states (NatureServe 2026). While *L. perennis* remains robust throughout much of its Southern range, populations in the Northeast and Midwest are often small and highly fragmented (Linton 2017; Michaels 2019). *Lupinus perennis* is a keystone species for pine barren ecosystems because it hosts more than a dozen insect pollinators, including the federally endangered Karner Blue and at-risk Frosted Elfin butterflies (Smallidge 1996; USFWS 2018; Ragan 2019). The species is therefore critical to the restoration of pine barren ecosystems.

A commonly cited gap in practitioner ability to restore pine barren ecosystems is the absence of genetic data for *L. perennis* populations. Microsatellite markers were first developed in the species to compare population size to inbreeding depression in paired Michigan and Ohio populations (Shi 2004). Michaels et al., 2008 reports no link between continuous population size (ranging from 149–3,264 individuals) or size category and inbreeding depression using these microsatellites. The same microsatellites were used in 2025 to assess population structure of the species across the Midwest and Northeast, which also found no relationship between population size and genetic diversity (Petitta et al. 2025).

Microsatellites are an inexpensive genetic method that have been used in conservation for decades, but their results can be imprecise with limited applicability to conservation planning (Ouborg et al. 2010). Whole genome techniques, especially those with a reference genome, allow investigation of population-level genetics with greater certainty and application to conservation (Nielsen et al. 2020; Formenti et al. 2022). At this point, no whole genome analyses have been published on *L. perennis*. Whole genome data can be particularly powerful in planning and assessing the effects of species translocations, which are a common part of ecosystem restoration interventions (Maschinski & Albrecht 2017).

Additionally, despite increasing cost-effectiveness of sequencing techniques, a wide gap remains in the implementation of genetic recommendations in conservation (R. Taylor et al. 2017; Schmidt et al. 2023). Persistence of the conservation genetics gap over the last decade is in part because of the deeply rooted differences in structure and incentive systems between management and research organizations (Shafer et al. 2015). The rapid evolution of conservation genomics has also created uncertainty in the reliability of untested methods and challenges resource-limited organizations to keep pace with a constantly changing field (Allendorf 2025). Conversely, policy and practice tend to adapt slowly and cautiously, which introduces further uncertainty in how genetic diversity management can and should be applied under outdated statutes (Sandström et al. 2019; Klütsch & Laikre 2021). When genetic data are applied to conservation challenges, the decision-making process behind these projects is often obscured or left unpublished, limiting contributions to the broader community (Garner et al. 2016). To implement conservation genetics into *L. perennis* restoration plans, it is therefore necessary to both understand the existing barriers for genetic-based recommendations in pine barren management and to generate whole genome data that will directly address practitioner concerns.

To characterize the potential conservation genetics gap in *L. perennis* restorations, we first assess knowledge, perception, and experience with genetics in the professional pine barren conservation community. We then apply a reduced representation whole genome sequencing technique to investigate the relationship among *L. perennis* populations in the Northeast, which represents the northern and eastern range edge of the species and has several organizations planning or implementing pine barren restorations. We retest the findings Michaels et al. (2008) and Petitta et al. (2025) reported to determine if there is a relationship between *L. perennis* population size and genetic diversity across a greater range of population sizes with whole genome data. Then, focusing on questions identified by Northeast *L. perennis* managers, we use our genomic data to model the minimum viable population size for the species and simulate effects of different potential seed sources on genetic diversity of newly founded subpopulations. To determine the best approach to closing the conservation genetics gap with our collaborators, we finally analyze the pine barren decision-making network to better understand how to communicate our codesigned results and recommendations to the community.

## Methods

### Pine Barren Management Survey Design

We developed an online survey administered by Qualtrics software (Qualtrics, Provo, UT) with twenty-one questions, including two mandatory and three response-dependent questions. The first question, asking for consent to participate in the research study, immediately concluded the survey if the ‘No’ option was selected. Five questions collected demographic information, including the respondents’ employers, job title, education, and geographic region. The remaining non-response dependent questions addressed two topics: 1) Perception and experience with genetics and 2) barriers to implementing conservation genetics. Response options to these topic questions included a five-point Likert scale (Likert 1932) (N = 6), multiple choice (N = 3), Yes/No (N = 2), and ranking (N = 1). Where appropriate, respondents were provided with opportunities to provide non-mandatory comments or to add their own options to questions. To decrease survey dropout rates, we conserved syntax across questions where possible and modeled our questions and response options on a successful survey of conservation professionals (R. Taylor et al. 2017).

The three response-dependent questions were only provided to individuals who indicated they worked in the Northeast of the United States. These questions aimed to investigate the conservation decision-making networks in this region while expanding the population of surveyed individuals through snowballing (Coleman 1958; Reed et al. 2009). We provided a list of regional organizations involved in these decision-making processes and asked respondents how often they were in communication on a Likert scale. Respondents were then asked if they communicated with organizations not on this list, and if so, to provide the names of these organizations and the frequency of communication on a Likert scale. All responses were anonymous and organizations were assigned a code to prevent matching responses to employees. For survey design, see supplementary materials.

This survey was approved as exempt from review by the University of Massachusetts Boston Institutional Review Board (#4086) on April 28, 2025. We began by identifying extant pine barrens in the Northeast and determining which policymaking bodies and management groups were responsible for their conservation. We then communicated directly with these organizations to determine which individuals had decision-making power. We began by distributing the survey to this set of practitioners directly responsible for managing pine barren ecosystems in the Northeast. We then sent the survey to organizations identified by respondents as involved in the pine barren conservation network. We repeated this process in successive rounds, sending the survey to new organizations identified in previous responses. Collection of survey data ran for one year beginning on May 12, 2025.

### Quantitative Survey Analysis

We investigated patterns in responses by calculating mean Likert scores and percentages from vote counts of multiple choice and yes/no answers. We conducted these and subsequent analyses in R (v. 4.4.1) using tidyverse (v. 2.0) packages (Wickham et al. 2019; R Core Team 2024).

Using the three response-dependent questions from our survey, we reconstructed the communication network for pine barren conservation in the Northeast. We used the R package igraph (v. 2.1.1) to calculate network statistics and ggraph (v. 2.2.2) to visualize the network (Csárdi & Nepusz 2006; Pedersen 2025). We finally created a genetics knowledge map by mapping data from survey responses as node attributes in the network graph.

### Genetic Material Sampling

We sampled leaves from *Lupinus perennis* populations across the Northeast United States in the 2022 growing season and worked with collaborators to obtain samples from the Midwest, Mid-Atlantic, and Florida (Figure 1A). We also obtained leaf samples from a Native Plant Trust seed bank germination test on a seed bank accession of the sole remaining Vermont population (VT1) collected in 1992. Sampling efforts resulted in a dataset of fourteen populations concentrated in the Northeast United States. Sites in the Midwest, Mid-Atlantic, and Southeast are all under conservation protection and are considered outgroups to the Northeast dataset. We surveyed each site to delineate *L. perennis* subpopulations, in consultation with conservation managers when possible. The number of plant samples collected varied based on population size and permitting requirements, with plants sampled haphazardly along transects running through the approximate center of selected subpopulations.

**Figure 1.**
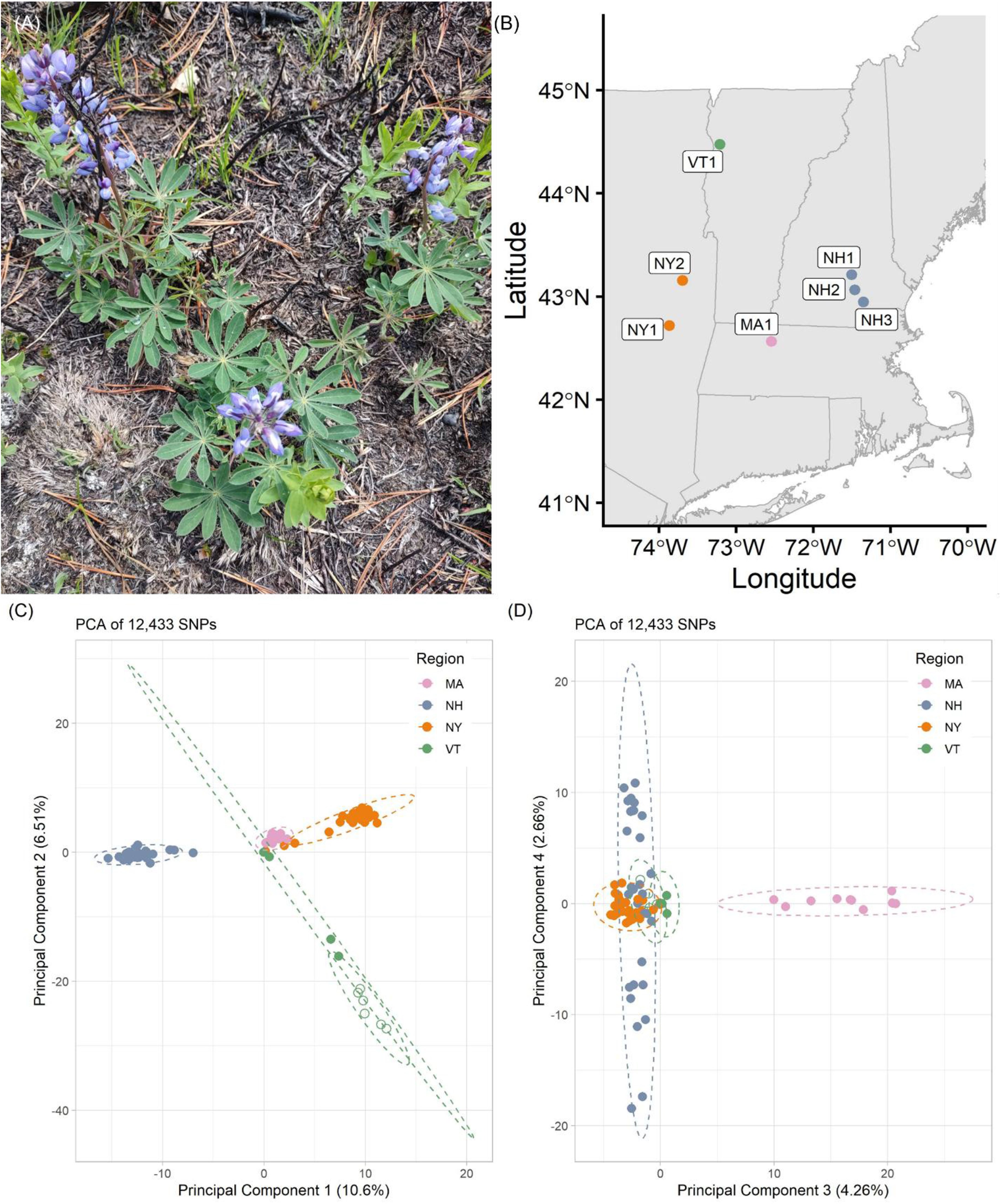
*Lupinus perennis* in the Northeast United States (A) A representative photo of a flowering *Lupinus perennis* individual in their third or fourth growing season. (B) A map of the populations sampled for genetic material in the Northeast U.S. Points on the map have been generalized and obscured to protect location information of protected populations. (C) Axis 1 and 2 of a principal components analysis calculated on 83 *L. perennis* samples from fourteen geographic locations and one seed bank accession based on 12,433 biallelic SNPs. (D) Axis 3 and 4 of the same PCA. Eigenvectors were calculated with *adegenet* with ellipses to estimate structure. Shape fill indicates contemporary (filled) versus historical (unfilled) collections from Vermont.

Samples from Florida were collected by Geena Hill with guidance from Jaret Daniels, while Midwest and Midatlantic sample collection was overseen by Isabella Petitta. Some samples were analyzed twice, once using our whole genome method and once using microsatellite loci (see Petitta et al. 2025). These included samples from populations I prepared (FL1, MA1, NH1, FL1) and samples from populations Petitta et al. 2025 prepared (IN2, MD2, MI2, PA1S) along with the NY1 population which both teams sampled and analyzed independently.

### Library Preparation and Sequencing

We generated Genotype-by-Sequencing (GBS) data for wild collected samples of *L. perennis* using a double digest Restriction-site Associated DNA approach (Poland & Rife 2012). We extracted DNA from leaflet tissue using a Qiagen DNeasy Plant Pro Kit (Cat. #69204). We next normalized concentrations and digested the extracts with MluC-I (New England Biolabs Cat. #R0538) and Msp-I (New England Biolabs Cat. #R0106) restriction enzymes. We then ligated custom forward and Y-shaped reverse adaptors (synthesized by Integrated DNA Technologies) corresponding to the digest cut sites with T4 DNA polymerase (New England Biolabs Cat. #M0202). Adaptors are designed to preferentially amplify sites that begin with the infrequent Msp-I cut site. We then amplified and added Illumina I5 and I7 adapters with barcoded primers (Integrated DNA Technologies XGen UDI Primer Plates: Plate 1, 8nt: product# 10005922; Plate 2, 8nt: product# 10009816). We carried out amplification for fourteen cycles with the Kapa HIFI HotStart Readymix (Roche Diagnostics Cat. #07958927001). Each sample was then quantified by High Sensitivity Qubit (Thermo Fisher Cat. #Q32854), renormalized, and pooled. The library was size selected using SPRI beads to a fragment size of 300–1000 bps. We SPRI bead cleaned and re-concentrated samples after ligation and amplification steps. Genewiz Azenta (Genewiz, South Plainfield, New Jersey) sequenced 150 bp paired-end fragments on two Illumina Novaseq X Plus lanes with 10% PhiX spike-in (see supplementary materials for adaptor sequences and full protocol).

### Population Genetic Analysis

We performed sequence quality control checks using fastqc and trimmed Illumina adaptors using trimmomatic (v. 0.39) (Andrews 2010; Bolger et al. 2014). We then aligned the paired end libraries against the *L. perennis* reference genome (So 2024) using bwa (v. 0.7.17) (Li & Durbin 2009). We then used samtools (v. 1.19.2) to filter alignments with qualities lower than 20 (Danecek et al. 2021). For population genetic analyses, we used the STACKS 2 (v. 2.68) ref_map.pl script to call SNPs (Rivera-Colón & Catchen 2022). We used the default value for minimum stack depth (-m 3) and tested distance between stacks within (-M) and between populations (-n) from 1 to 10. The default parameters (M=n=3) performed best on an r80 test and were used to generate a VCF file filtered for biallelic SNPs with a maximum 80% missingness and a minimum minor allele frequency of 1% (Paris et al. 2017). We applied the same filtering to produce VCFs of all populations and of only Northeast populations. Samples from the Vermont 1992 seed bank accession were included as a separate population from the 2022 sampling.

We examined genetic population structure by constructing a PCA using the R package adegenet (v. 2.1.10) (Jombart 2008). Missing SNP calls were replaced with the mean of the missing sample’s population for that locus. We constructed a PCA of samples from all Northeast populations (Figure 1B). We then used poppr (v. 2.9.8) to calculate population diversity summary statistics and hierfstat (v. 0.5.11) to estimate F-statistics using the function “genet.dist()” and heterozygosity using the function “basic.stats()” (Goudet 2005; Kamvar et al. 2014). Excluding the 1992 seed bank accession, we assessed isolation by distance by comparing pairwise matrices of Weir and Cockeram’s F_ST_ and straight-line distance between populations using a Mantel test from the vegan (v. 2.6-8) package (Weir & Cockerham 1984; Jari et al. 2001). We also constructed linear models to predict the relationship between population area as a predictor of inbreeding and observed heterozygosity (F_IS_ ∼ area; H_O_ ∼ area), which we assessed using the performance (v. 0.12.3) and broom (v. 1.0.6) packages (Lüdecke et al. 2021; Robinson et al. 2025).

### Minimum Viable Population Size Analysis

To inform ongoing conservation efforts to protect *L. perennis*, we estimated the minimum viable population (MVP) size for the species using Vortex (v. 10.10.0) to stochastically model the effects of periodic catastrophes on extinction probability (Lacy & Pollak 2025; Lacy et al. 2025). Our Vortex model of MVP is parameterized with existing literature on *L. perennis*, much of which has already been summarized and reanalyzed by the USDA and US FWS for conservation purposes (Meyer 2006; USFWS 2018; Ragan 2019; Petitta et al. 2024). See Supplementary Materials for full notes on simulation inputs.

When assessing populations for extirpation risk, conservative MVP estimates should account for future variation in catastrophe rates. According to subpopulation monitoring data, a general estimate is that *L. perennis* experiences a 50% die-off once per generation (or a 5% chance per year), which aligns with estimates in vertebrate populations (Reed et al. 2003). However, catastrophes are unpredictable and will vary in both space and time. We define MVP size as the initial population size conferring 99% probability of persistence for 40 generations, or 140 years in the case of *L. perennis* (Flather et al. 2011). We tested every combination of initial population size from 1–250 incrementing by one and catastrophe frequency from 0–10 incrementing by a half percent point. Carrying capacity was set to the same value as initial population size for all simulations. The model ran 300 iterations for each pair of parameter values over five independent replicates for a total of 7.5 million populations over 140 simulated years. We then summarized the sensitivity test data to determine the minimum initial population size resulting in less than 1% probability of extinction for each catastrophe frequency in each replicate. We constructed models of the relationship between MVP and probability of catastrophe with modelr (v. 1.11) and tested if there was greater support for a linear or quadratic relationship using AICcmodavg (v. 2.3-4) (Mazerolle 2023; Wickham 2023). We assessed fit of the better-supported model using performance and calculated coefficients with broom.

### Founded Population Simulations

We modeled the genetic effects of founding new populations at the MVP size using seed from existing Northeast *L. perennis* populations. We categorized existing potential donor populations based on census size as small (dozens of plants: MA1, NH3), medium (hundreds of plants: NH2, NY2), and large (tens of thousands of plants: NH1, NY1). Pairings between NH sites as one donor and NY/MA sites as the other had similar pairwise F_ST_ values, providing two replicates of each seed source size category where genetic differentiation between populations is controlled. We provided Vortex with the measured allele frequencies of the 12,433 SNPs variant among the Northeast population samples to calculate average effective population size (*Ne*) over 140-year simulations. Each seed source contributed half of the MVP size to the founded population with alleles drawn from each source’s specific set of calculated frequencies. We set the global catastrophe parameter, which affects each population equally, at a 5% frequency per year (Reed et al. 2003). Founded populations from each seed source pairing were simulated over six replicates with 250 iterations each. We also simulated populations founded from a single seed source at the same MVP size for each of the six potential donor sites. We constructed gamma generalized linear models to assess the relationship between *Ne* and number of seed sources (*Ne* ∼ N seed sources) and seed source size (*Ne* ∼ size category) with glmmTMB (v. 1.9.19) (Brooks et al. 2017). We compared means of *Ne* over the 140 year period for each replicate simulation using a tukey test in emmeans (v. 1.10.4) to test if founding populations from two seed sources resulted in higher average *Ne* and if source population size influences *Ne* (Lenth, 2024).

## Results

### Defining the Pine Barren Conservation Genetics Gap

We received nineteen responses from individuals consenting to participate in our pine barren conservation research survey for a 37% response rate at the organizational level. These individuals were associated with fourteen organizations: five nonprofits, eleven state agencies, and three federal departments. All respondents held a professional conservation biology position and had a four-year college or graduate degree. All landowners, managers, and permitting agencies that oversee the *L. perennis* populations we sampled responded to the survey.

We first assessed survey responses on the use of genetic data to inform conservation challenges. Respondents were asked to score how useful genetic data is to address taxonomy, captive breeding, inbreeding, restorations, translocations, and population connectivity. Mean Likert scale values for all six categories ranged from 3.84 to 4.26 (n = 19), corresponding to an average response of “Very useful” (Figure 2A). Opinions on using genetic data to address population connectivity were the most mixed (sd = 0.958) while restorations had the lowest spread of responses (sd = 0.667). The consistent high Likert scoring suggests professionals understand that genetics can be applied to a range of conservation problems and that doing so would be useful. We then aimed to better characterize conservation professionals’ experience with genetics by asking them which of these categories they have applied genetics to in their work (Figure 2B). Nearly one-third of the response pool (n = 6) had never applied genetics to these topics in their past work. The most commonly cited uses of genetic data were to address species or population restorations (n = 9, 47.4%) and taxonomy (n = 8, 42.1%). The least cited genetic application was to captive breeding (n = 2, 10.5%), which may reflect the relative rarity of this conservation intervention compared to the other categories. These responses highlight the conservation genetics implementation gap and suggest that barriers exist beyond professional knowledge of genetic method use.

**Figure 2.**
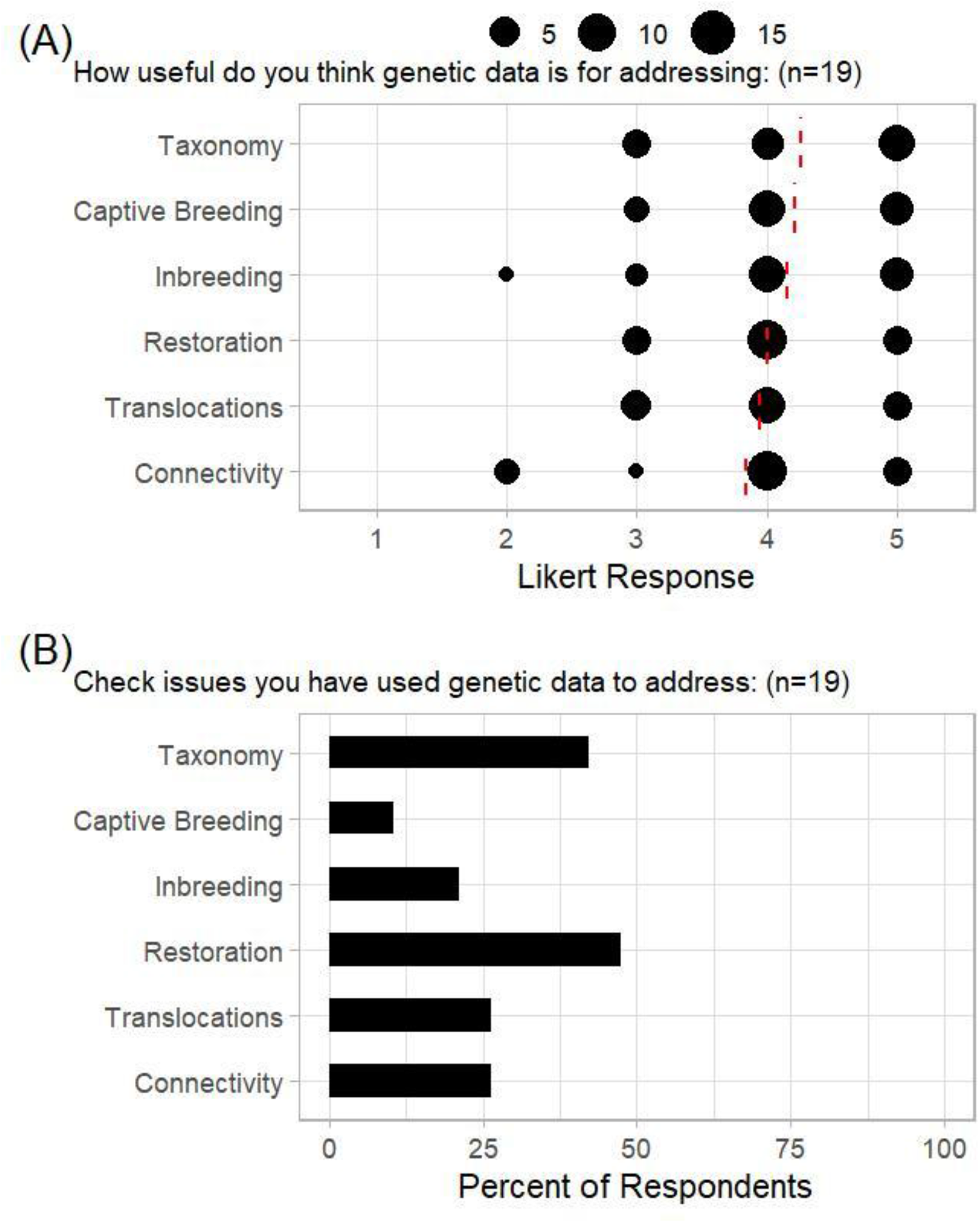
Perceptions of the usefulness of genetic data among conservation professionals Survey participants were asked how useful genetic data is to address common topics in conservation biology and if they have made use of genetic data for past interventions (*n* = 19 for all questions). (A) Responses on usefulness are scored on a Likert scale (1 = Not at all useful, 5 = Extremely useful) with lines indicating the mean response. (B) Proportion of respondents who indicated past use of genetic data for each challenge category. Respondents were allowed to select multiple categories from the list or indicate that they had not used genetic data for any of these purposes.

We then asked a series of questions to identify barriers in the implementation of conservation genetics (Figure 3A). Respondents indicated that they believe genetic diversity is very important to population persistence (mean Likert score = 4.21, n = 19) and that they are somewhat to very likely to use genetic resources for future projects if available (mean Likert score = 3.58, n = 19). Respondents were split on how often they take genetics into consideration for their management plans, with most indicating they do so sometimes or most of the time (mean Likert score = 3.12, n = 17). However, respondents were uncertain if they had the knowledge to personally use genetic recommendations (mean Likert score = 2.89, n = 19), and only one fifth (n = 4) indicated that they had someone on their team with genetic experience.

**Figure 3.**
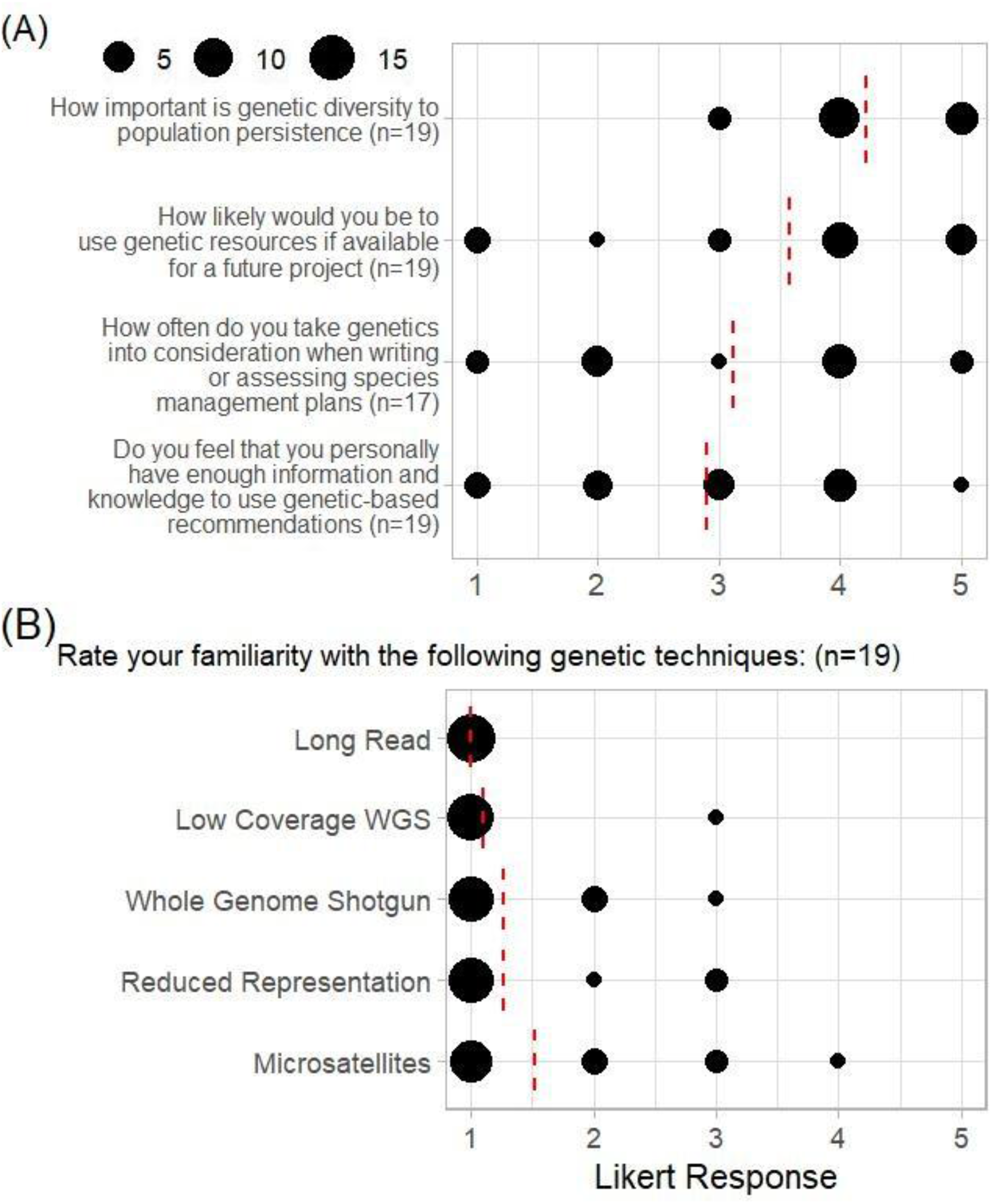
Evidence of a conservation genetic gap in Northeast pine barrens (A) Survey participants were asked a series of questions to gauge their perception of conservation genetics and their ability to use genetic recommendations with Likert scale responses (1 = low; 5 = high). (B) Likert scale responses to questions on respondents’ familiarity with genetic sequencing techniques. Red dashed lines indicate the mean response value.

The genetic knowledge gap is further evident when respondents were asked about their familiarity with five common genetic sequencing techniques (microsatellites, reduced representation, whole genome shotgun, low coverage whole genome shotgun, and long reads). Mean Likert scale responses ranged from 1 to 1.53, indicating that these conservation professionals were “Not at all familiar” to “Somewhat familiar” with the array of genetic techniques (Figure 3B). Respondents were most familiar with microsatellites, which have been a mainstay in conservation since the 1990s, while every respondent indicated they were not at all familiar with long read sequencing, the newest technology. When asked to identify reasons why they did not use genetics in past projects, 80% of respondents (n = 12) indicated that they did not have a contact or staff who could perform the work while 53% (n = 8) indicated they had insufficient funds for genetic testing. Only two respondents indicated that they did not know genetics were an option for their project. Our survey of well-educated, experienced conservation professionals illustrates that the conservation genetics gap is not one of willingness or knowledge, but one of resources. We fill this gap for *L. perennis* restoration projects by funding, producing, and analyzing genetic data for managed populations of the species in the Northeast.

### Lupinus perennis *Population Genomics*

We generated a dataset of 142 successfully sequenced plant samples from fourteen geographic populations and one paired seedbank accession. After filtering on missingness and minor allele frequency, we had 40,104 biallelic SNPs variable among all samples, with 12,433 variable among the 83 plants sampled from Northeast populations (Figure 1B).

The genetic Principal Components Analysis (PCA) of the Northeast samples indicates separation along PC1 between the samples from New Hampshire and those from New York and 1992 Vermont, explaining 10.6% of variance in the data (Figure 1C). PC2 explains 6.51% of variance and is primarily driven by a temporal difference in sampling with those collected in 2022 differentiating from the Vermont seed bank accession collected in 1992. Explaining 4.26% variation, PC3 is the last component to show differentiation among populations, in this case driven by differences between Massachusetts and all other samples (Figure 1D). An additional 76 principal components have eigenvalues greater than one, but visual inspection of these axes indicates variation driven by diversity within rather than among populations, as seen in PC4.

The Northeast populations exhibit isolation by distance (*⍴* = 0.785, *p* = 0.001 for a Mantel test of F_ST_ ∼ distance). Pairwise genetic differentiation is low among these populations, from a minimum F_ST_ of 0.005 between NY1 and NY2 to a maximum F_ST_ of 0.037 between VT1-22 and both NH2 and NH3. Similarity between the New York populations is expected as NY1 provided seeds for major restorations of the NY2 population, while geographic features including the Green Mountains may shape the separation between the New Hampshire and Vermont populations.

Some population genetic metrics appear less shaped by distance and more by population size (Table 1). Population diversity indices are lowest in the smallest VT1-22 population (Shannon-Weiner’s H: 1.386, Simpson’s λ: 0.750) and highest in the largest NY1 population (Shannon-Weiner’s H: 2.996, Simpson’s λ: 0.950). Heterozygosity estimates, however, do not follow this pattern. The Southeast United States is thought to be the origin of diversity for this species, which is supported by expected and observed heterozygosities an order of magnitude larger in the Florida populations than in any others sampled. Within the Northeast, the VT1-22 population again has the lowest diversity values (*H_O_* = 0.016, *H_E_* = 0.017), while all of the NH, MA, and NY populations have similar heterozygosities (*H_O_* = 0.032–0.035, *H_E_* = 0.036–0.038). A linear model predicting heterozygosity from population area (*H_O_* ∼ Area in thousands of acres) indicates no relationship in Northeast populations (Estimate = 0.003, *p* = 0.357). This suggests that while the largest and smallest heterozygosities align with size ranking, the two are not consistently predictable across the range of population sizes in this dataset.

**Table 1.** *Lupinus perennis* Population Genetic Statistics. Summary statistics calculated with *poppr* include sample size (*N*), Shannon-Weiner Diversity Index (*H*), Simpson’s Diversity Index (*Lambda*), and number of private alleles (*A_P_*). Statistics calculated by *hierfstat* include observed (*H_O_*) and expected (*H_E_*) heterozygosity and inbreeding (*F_IS_*) with bootstrapped 95% confidence intervals.

| Population | $N$ | $H$ | $Lambda$ | $A_P$ | $H_O$ | $H_E$ | $F_{IS}$ | $F_{IS}$ 95% CI |
| --- | --- | --- | --- | --- | --- | --- | --- | --- |
| FL1 | 8 | 2.079 | 0.875 | 2457 | 0.142 | 0.149 | 0.045 | (0.038, 0.052) |
| FL2 | 10 | 2.303 | 0.900 | 5213 | 0.116 | 0.142 | 0.185 | (0.179, 0.191) |
| FL3 | 11 | 2.398 | 0.909 | 4236 | 0.123 | 0.146 | 0.154 | (0.147, 0.16) |
| IN2 | 5 | 1.609 | 0.800 | 271 | 0.024 | 0.029 | 0.153 | (0.129, 0.177) |
| MA1 | 10 | 2.303 | 0.900 | 306 | 0.033 | 0.036 | 0.066 | (0.054, 0.079) |
| MD2 | 8 | 2.079 | 0.875 | 141 | 0.029 | 0.040 | 0.239 | (0.218, 0.261) |
| MI2 | 8 | 2.079 | 0.875 | 222 | 0.021 | 0.028 | 0.250 | (0.23, 0.27) |
| NH1 | 12 | 2.485 | 0.917 | 251 | 0.034 | 0.037 | 0.083 | (0.073, 0.094) |
| NH2 | 9 | 2.197 | 0.889 | 141 | 0.032 | 0.037 | 0.116 | (0.105, 0.127) |
| NH3 | 9 | 2.197 | 0.889 | 182 | 0.033 | 0.036 | 0.065 | (0.053, 0.078) |
| NY1 | 20 | 2.996 | 0.950 | 554 | 0.034 | 0.038 | 0.101 | (0.091, 0.11) |
| NY2 | 13 | 2.565 | 0.923 | 89 | 0.035 | 0.038 | 0.086 | (0.077, 0.095) |
| PA1S | 9 | 2.197 | 0.889 | 113 | 0.028 | 0.035 | 0.157 | (0.134, 0.18) |
| VT1-22 | 4 | 1.386 | 0.750 | 18 | 0.016 | 0.017 | -0.128 | (-0.204, -0.052) |
| VT1-92 | 6 | 1.792 | 0.833 | 611 | 0.026 | 0.031 | 0.157 | (0.139, 0.175) |

We calculate inbreeding as F_IS_ with bootstraping to estimate 95% confidence intervals (Table 1). Inbreeding in one population, VT1-22, is consistently negative (*F_IS_* 95% CI: -0.204, -0.052), while its seed bank accession VT1-92 is only slightly higher than the other Northeast populations (*F_IS_* 95% CI: 0.139, 0.175). The highest inbreeding levels were observed in the MI2 (*F_IS_* 95% CI: 0.23, 0.27) and MD2 (*F_IS_* 95% CI: 0.218, 0.261) populations. Notably, all but one of the Midwest and Florida populations have higher inbreeding estimates than all of those in the Northeast. A linear model predicting inbreeding from population area (*F_IS_* ∼ Area in thousands of acres) again indicates no relationship (Estimate = 0.028, *p* = 0.416). These results suggest that inbreeding estimates are not sufficient for categorizing at-risk populations, necessitating an alternate approach.

### Minimum Viable Population Analysis

As *L. perennis* declines across its natural range, a key value in assessing risk of extirpation is the Minimum Viable Population (MVP) size. Given similarities in genetic inbreeding across the Northeast populations, this non-genetic method may be of use in determining which populations are most in need of intervention. Our modeled Vortex sensitivity tests follow a logistic curve, with near 100% extirpation at low initial sizes and near 0% at high initial sizes (Supplementary Figure 1A). As expected, the rate of catastrophe per year (defined as a 50% mortality event) greatly impacts probability of extirpation, and therefore MVP (Supplementary Figure 1B). As the effects of climate change worsen and catastrophe events become more common, the necessary MVP to prevent extirpation will also increase. Our model of MVP predicted by probability of catastrophe (MVP ∼ Catastrophe%) is best fit as a raw quadratic polynomial relationship (y = 0.366x^2^ + 3.43x + 41.7; Adjusted r^2^ = 0.76). The model predicts that the upper confidence interval of estimated MVP at a 5% catastrophe rate per year is 71, while at 10% it is 113.

### Simulating Seed Source Effects on Founded Populations

We first modeled the effect of founding an MVP sized population (71 individuals) with one or two seed sources, using calculated allele frequencies from Northeast populations to estimate mean *Ne* over 40 generations with a 5% annual rate of catastrophe. There is no detectable difference in average *Ne* between populations founded from one or two seed sources (Estimate: -0.121, 95% CI: -0.100-0.341, *p* = 0.28 for a means test based on a gamma GLM). Within the populations founded from a single seed source, there is no effect (Kruskal-Wallis *p* = 0.97) of the size of the source population on average *Ne* (Figure 4A). This result is supported by pairwise means tests between small, medium, and large population effects on average *Ne* based on a Gamma model, which estimated all effect size confidence intervals overlapping zero. This result is replicated in the populations founded from two seed sources: no effect of population size pairings on average *Ne* (Kruskal-Wallis *p* = 0.604) and all effect size confidence intervals overlapping zero (Figure 4B).

**Figure 4.**
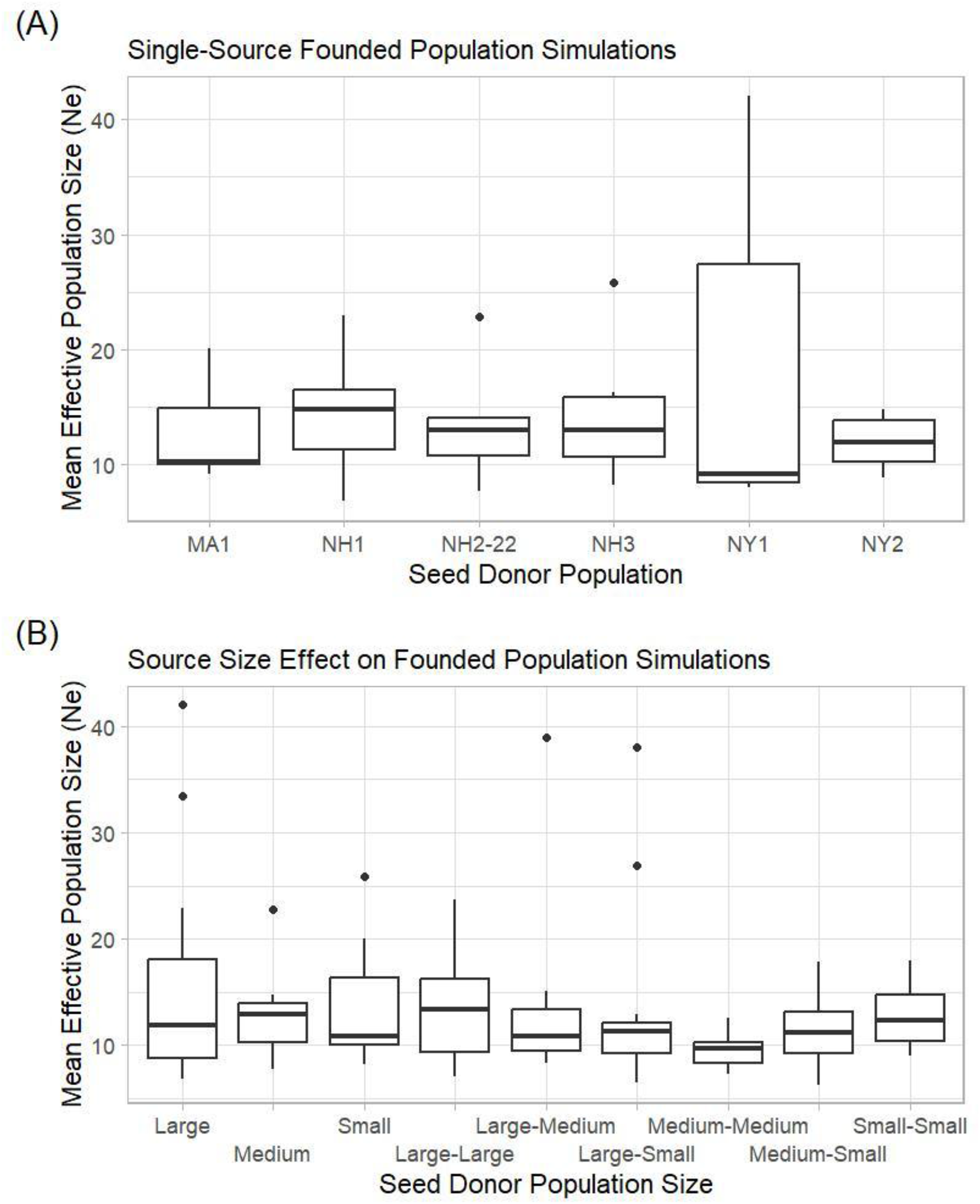
*Ne* of simulated populations from single or two seed sources (A) *Vortex* simulated effective population size (*Ne*) of founded populations from each potential *L. perennis* seed source in the Northeast. (B) *Vortex* simulations of founded *L. perennis* populations from two seed sources, grouped by size category. Each simulation was rerun in six replicates with 250 iterations each to calculate mean *Ne* over a 140 year (40 generation) period with a 5% annual rate of catastrophe. All catastrophes are global and affect each population within the iteration at the same time. No average *Ne* group means differ among single seed sources, paired seed sources, or between all combined single and paired seed sources.

Parsing the simulations year-by-year shows high and unpredictable variance in *Ne* within each founded population, with a nearly negligible effect of year on *Ne* (Estimate = -0.0003, *p* = 0.05) even as populations are generally expected to lose genetic diversity over time due to drift. These results are potentially derived from the overall similarity in heterozygosity, genetic diversity, and inbreeding measured across the Northeast populations on which these simulations are based. While these populations show minor genetic differentiation, stochastic events may obscure seed source effects on *Ne*.

### Pine Barren Decision-Making Network

Implementing interventions that involve seeds of multiple sources requires coordination between conservation stakeholders, including population managers, policymakers, and seed providers. Using a final set of questions from our survey, we mapped the decision-making network in pine barren ecosystems in the Northeast. Our network has significant representation in the state, federal, and nonprofit spheres but lacks any privately owned entities. Respondents (n = 15) identified four federal, eleven state, and eighteen nonprofit organizations as part of the Northeast pine barren conservation decision-making network (Supplementary Figure 2A). Thirty percent (n = 10) of these organizations directly manage a pine barren ecosystem or species. Across the full network, respondents reported 115 decision-making connections between organizations with an average communication frequency of “3–5 times per year” (mean Likert score = 3.03, sd = 0.976). Organizations in the network had a high variance in degree (mean = 6.97, sd = 7.00) and betweenness (mean = 9.24, sd = 20.0) suggesting that the network is dominated by several well-connected nodes. These well-connected organizations drive a short distance between nodes (mean = 1.98) despite the high number of relatively unconnected edge groups. The eight central organizations to the network are representative of the whole, including one federal, three state, and four nonprofit organizations.

Of the fourteen organizations we received survey responses from, eight identified as pine barren ecosystem or species managers. This subset of nodes is important not just because of their influence on decision-making, but also because they carry out the decisions themselves (Supplementary Figure 2B). Management organizations reported a total of 31 connections among themselves, but with less variance in node degree (mean = 7.4, sd = 3.84) and betweenness (mean = 5.5, sd = 6.08) than the larger network. The distance between manager nodes is also similar to the whole network (mean = 1.75) but this subset of organizations is less dominated by a small number of well-connected nodes. Reported communication frequency was the same in the manager network as the full set, corresponding to “3–5 times per year (mean Likert score = 3.02, sd = 1.04). Finally, knowledge mapping of responses to the earlier question on ability to use genetic recommendations reveals that management organizations were unsure of their ability to implement genetics (mean = 3.37, sd = 1.06, n = 10). However, responses indicate that there are well-connected members of the network who feel they have the knowledge and resources to do so.

## Discussion

Conservation decision-making is a complex process relying on the perceptions, experience, and regulatory restraints of specific management organizations and their partners. Barriers to incorporating genetics into these processes are important to understand, particularly as sequencing has become cheaper over the last decade (Shafer et al. 2015). The conservation professionals in our study expressed knowledge of the usefulness of genetics to plan, implement, and monitor their interventions. They also showed awareness of the benefits of genetic data, even without understanding specific genotyping or sequencing techniques. This technical knowledge gap has not prevented these same managers from using genetic recommendations in the past, with nearly half reporting use for restoration and taxonomy. Our network analysis also shows that pine barren managers are not an insular community. They are well connected with one another and to organizations outside of direct management, giving access to genetic knowledge and expertise even to those that do not possess it internally.

The managers to whom we first sent our survey were those that we had already contacted to gain permission for *L. perennis* genetic material sampling. While these organizations represent the majority of *L. perennis* populations in the Northeast United States, their managers may represent a skewed demographic who were already willing to participate in genetic research. Our snowballing technique for survey distribution may mitigate this effect across the larger network, but these results should be viewed through the lens of some self-selection bias. Similarly, this decision-making network likely appears more connected than managers are in reality, as most respondents were from organizations that managers identified as decision-making partners. Isolated conservation organizations would therefore never be identified using our distribution approach, inflating the measured connectivity of our analyzed network. However, based on our experience with the extant pine barrens in the Northeast United States, we believe that the most influential organizations for management of these ecosystems regionally are represented in our study.

The genetic methods, questions, and analysis approach we use in our research were all designed with the goal of directly answering conservation managers’ questions about how at-risk their populations were and where they should source seeds for restorations. Our initial hypothesis that population size would correlate with genetic diversity of both extant and simulated populations was unsupported by our data, just as it was unsupported in the Midwest and Midatlantic *L. perennis* populations Michaels et al. (2008) and Petitta et al. (2025) analyzed with microsatellites. Our results show an unexpectedly positive picture for many of the sampled *L. perennis* populations, with the notable exception of VT1-22, our smallest sampled population. Overall, Northeast *L. perennis* populations have similar levels of genetic diversity, heterozygosity, and inbreeding despite orders of magnitude in habitat area and census size variation. A continual challenge in conservation genetics is identifying reference populations that have not themselves been shaped by ecosystem destruction and potential bottlenecking; pine barrens as an ecosystem are defined as much by their species as their rapid disappearance. Even the largest population in our study, the Albany Pine Bush (NY1), experienced rapid ecosystem destruction through the late 1900s before substantial conservation and restoration efforts shaped it into a preserve of more than 3,000 acres. Our results could be interpreted as a sign that genetic diversity in *L. perennis* has largely persevered, perhaps due to the plant’s perennial life history, obligate outcrossing reproduction, and robust natural seedbank. Alternatively, our results may show a species after region-wide bottlenecking with too little time for the trajectory of larger and smaller populations to measurably separate.

Our results partially match the patterns that Petitta et al. (2025) report from their microsatellite-based analysis of some of the same populations and samples. We both report that genetic diversity is highest in Florida and that populations show signs of genetic isolation by distance. Our population inbreeding estimates (F_IS_) differ, with our GBS method of 40,104 SNPS reporting low levels with high confidence compared to a high uncertainty result from their 25 microsatellites. We also find a clearer pattern in genetic differentiation, with our lowest F_ST_ value corresponding to NY1 and NY2 (seed from the former was recently used to restore the latter) and their lowest G_ST_ corresponding to NH1 and MD2 (two geographically distinct populations). Together, our primarily Northeastern focus and their Midatlantic focus provide a broader map of the state of *L. perennis* population genetics across its Northern range that will be useful in assessing and contextualizing risk in these populations.

### Management Recommendations

Recent standardization attempts define MVP as the initial size that results in less than one percent probability of extirpation over forty generations, which we calculate to be 71 *L. perennis* individuals at a 5% rate of catastrophe. At the time of sampling, two Northeast populations (MA1, VT1-22) are under this MVP cutoff. Both populations are actively monitored and are undergoing augmentation. One additional population (NH3) falls under the 113 MVP cutoff for a 10% catastrophe rate. This population grows on a roadside public right of way and is not currently under conservation protection. Our founded population simulations agree that genetic diversity (here modeled as *Ne*) of new *L. perennis* populations will not differ regardless of which or how many Northeast natural populations the seeds are sourced from. In part, this is a byproduct of the realistic stochasticity introduced by Vortex models, both in the process of allele inheritance and in which individuals die during catastrophes. Utilizing other stochastic models like SLiM or deterministic inheritance equations might provide yield different recommendations, but how these alternative approaches translate to observed restoration outcomes is unknown (Haller et al. 2026).

Applying our conservation genomic approach to ongoing restoration planning, we only exclude VT1-22 as a possible seed source, which would be difficult given the number of plants remaining in the population. Our genetic differentiation and isolation-by-distance results suggest that more distant populations will be less similar to a restoration site, but our PCA-based genetic structure analysis suggests that this explanation is overly simplistic. In the Northeast, genetic structure appears driven by differences among the New Hampshire and New York/Vermont populations and then between the Massachusetts population and the rest of the region. These patterns emerge despite approximately equal pairwise distances between VT1-22 and NY1, and VT1-22 and MA1. This suggests that geographic features, like the Green Mountains in Vermont, may act as a gene flow barrier across the region.

Taken together, our population genetic and structure results suggest that there is a promising opportunity for managers to engage in climate-smart seed sourcing for their restorations. Translocating seeds between populations that have shared evolutionary histories lowers the likelihood of genetic swamping and risk of maladaptation. Additionally, as a perennial species, translocated *L. perennis* seed must survive local conditions for several years before reproducing, further lowering these risks. Several tools exist to map locations that match an area’s future climate conditions, allowing managers to identify populations that could confer climate pre-adaptation if translocated (St.Clair et al. 2022). Currently, few studies exist that analyze the fitness effects of such translocations and none that measure genetic effects of climate-smart adaptation in plant species. By establishing a genetic data baseline for Northeast *L. perennis* and collaborating with the managers of these populations, we enable future studies of the fitness and genetic effects of climate-smart seed translocations.

This research has already shaped seed sourcing policy for ongoing restorations of the VT1 and NH1 *L. perennis* populations. The VT1 population is the last remaining in the state, numbering twenty-two plants in 2022 and twelve in 2024. A Native Plant Trust seed bank accession collected in 1992 from this population (VT1-92) is being grown and outplanted with the primary goal of preventing state extirpation of the species (G. Glynn, personal communication). Given the genetic diversity, private allele, and heterozygosity discrepancies between these two populations, managers hope that the VT1-92 seed bank will restore lost diversity to the extant population and improve fitness.

In New Hampshire, monitored *L. perennis* subpopulations at NH1 have declined by approximately 50% over the last twenty-five years, with an estimated 35,000–50,000 plants remaining (H. Holman, personal communication). While at little risk of extirpation, NH1 is too small to support a stable population of the federally endangered Karner Blue Butterfly. Managers are therefore augmenting the NH1 population with seed from NY1 (New Hampshire Fish and Game Department 2025). In addition to being the only population in the region capable of providing the amount of seed necessary for this augmentation, NY1 also matches future climate conditions expected at NH1. This seed translocation, using the genetic data presented here as a baseline, will directly assess the effects of a climate-smart seed translocation as a living experiment.

Testing these interventions with relatively low-risk populations provides an opportunity to develop translocation methods for future use with threatened species. Designing restorations to analyze genetic effects while still accomplishing conservation goals requires close collaboration with ecosystem managers and a deep understanding of their decision-making processes. We strongly recommend that managers undertaking restorations or translocations record the provenance of their seed, save genetic material for future testing, and incorporate basic fitness metrics (e.g. seed mass, number of flowers, etc.) into their monitoring plans. We also recommend that researchers looking to create actionable science begin by understanding the specific needs of their management partners prior to designing experiments (Cravens et al. 2026). An interdisciplinary approach to research codesign has the potential to broadly improve conservation biology and begin closing the conservation genetics gap one management network at a time.

## Acknowledgements

We thank collaborators Isabella Petitta for sharing DNA samples from four study sites and Cameron So, Dan Schoen, and Anna Hargreaves for sharing the unpublished draft *L. perennis* genome used in our analyses. Thank you to Albany Pine Bush, Florida Department of Agriculture and Consumer Services, Indiana DNR, MassWildlife, New Hampshire Fish and Game, New Hampshire Natural Heritage Bureau, New York State Department of Environmental Conservation, Nokuse Plantation, the U.S. Forest Service, Vermont Fish & Wildlife, and Wilton Wildlife Preserve for permitting genetic sampling of *L. perennis* on land they manage. Thanks also to Native Plant Trust for providing access to their *L. perennis* seed bank accession and for sharing monitoring data on extant populations.

Special thanks to all of the conservation professionals who shaped this research with invaluable insight into their management priorities and practice, including Heidi Holman, Grace Glynn, Michael Piantedosi, Jessamine Finch, Bob Popp, Neil Gifford, Kathleen O’Brien, and Chris Buelow. Additional thanks to the anonymous participants in our research survey and to the University of Massachusetts Boston Institutional Review Board.

This research was funded in part by the New England Botanical Society, the UMB Goranson Fund, and the UMB Transdisciplinary Dissertation Proposal Development Program. CKR was partially funded by a National Science Foundation Graduate Research Fellowship (Award #2235034) while working on this project.

The authors declare no conflicts of interest.

## Data Availability

Raw DNA sequencing files are openly available in NCBI’s SRA (PRJNA1516668 currently non-public). All genomic analysis scripts are archived at 10.5281/zenodo.22101713.

## Ethics Approval Statement

Our scientific survey was approved as exempt from review by the University of Massachusetts Boston Institutional Review Board (#4086) on April 28, 2025.

## Author Contribution Statement

CKR obtained funding for the study. CKR and GM designed the scientific survey and social network analysis. BM and CKR designed the genetic experiment. CKR conducted field work and all analyses. CKR and STG conducted molecular lab work. CKR wrote the first draft of the paper. All authors edited and approved the final draft of the manuscript.

---

**Supplementary Figure 1.**
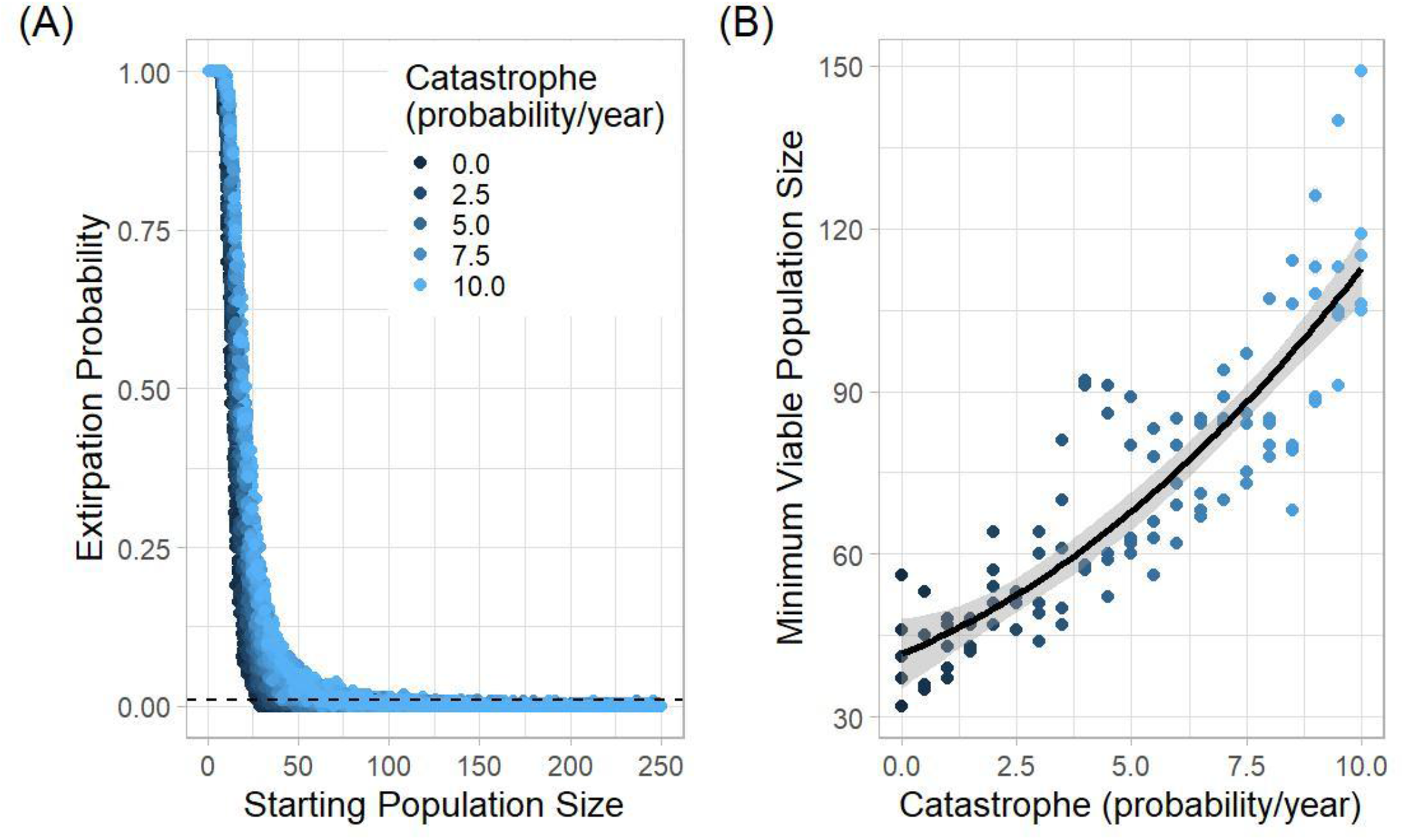
Minimum Viable Population simulation with variable catastrophe occurrence (A) Sensitivity test of the effects of initial population size and probability of catastrophe on population extirpation. *Vortex* simulated populations with factorial pairings of whole number population sizes from 1 to 250 individuals and each half percent catastrophe probabilities from 0 to 10. Each variable pair was run over five replicates with 300 iterations for 140 years (approximately 40 generations). (B) Modeled relationship of catastrophe probability as a continuous predictor of minimum viable population size with *modelr*. The best model as determined by *AICcmodavg* is a quadratic polynomial (y = 0.366x^2^ + 3.43x + 41.7; Adjusted *r^2^* = 0.76)

**Supplementary Figure 2.**
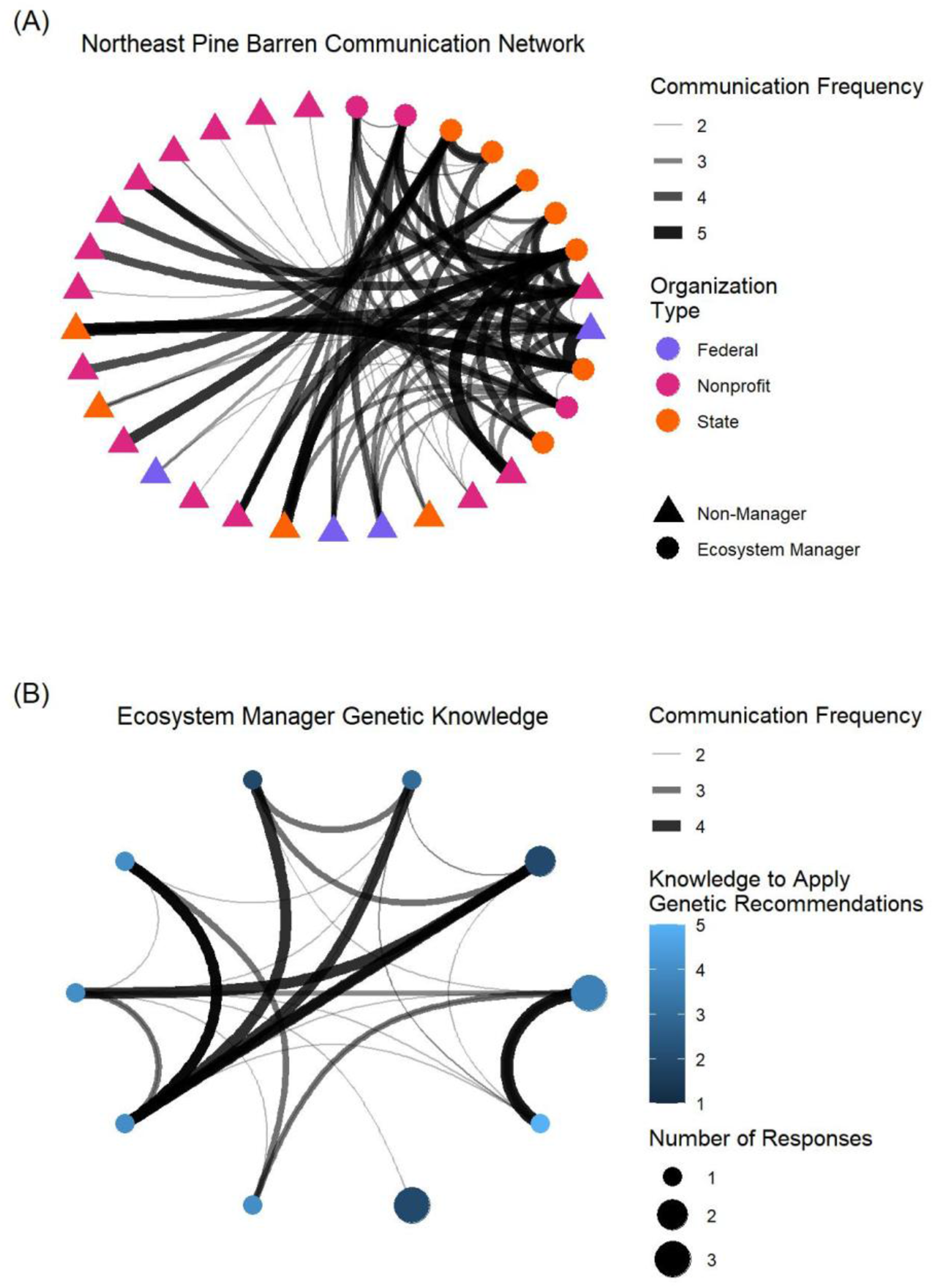
Pine barren conservation organization decision-making networks (A) A network graph of survey responses depicting communication among pine barren conservation organizations in the Northeast colored by organization type. Nodes represent organizations that we received survey responses from and those that respondents indicated were consulted during their conservation decision-making process. Respondents were asked to rate the frequency of communication with other organizations on a Likert scale, represented as width and opaqueness of connecting arcs. (B) The same network graph subset to include only organizations that directly manage pine barrens and their species. Nodes are sized based on the number of responses from each organization and are colored based on the average Likert scale response to a survey question asking managers to indicate their confidence in applying genetic recommendations to conservation.

## Literature Cited

1. Allendorf FW. 2025. Fifty Years of Conservation Genetics: A Personal Perspective. Molecular Ecology 34:e17705.

2. Andrews S. 2010. FastQC: A Quality Control Tool for High Throughput Sequence Data.

3. Bolger AM, Lohse M, Usadel B. 2014. Trimmomatic: a flexible trimmer for Illumina sequence data. Bioinformatics 30:2114–2120.

4. Brooks ME, Kristensen K, Van Benthem KJ, Magnusson A, Berg CW, Nielsen A, Skaug HJ, Mächler M, Bolker BM. 2017. glmmTMB balances speed and flexibility among packages for zero-inflated generalized linear mixed modeling.

5. Clark KS, N.; Gallagher, M. 2015. Fire Management and Carbon Sequestration in Pine Barren Forests. Journal of Sustainable Forestry 34:125–146.

6. Coleman JS. 1958. Relational Analysis: The Study of Social Organizations with Survey Methods. Human Organization 17:28–36.

7. Cravens AE, et al. 2026. Debunking the myth of the quintessential resource manager: Precision in actionable science. Conservation Science and Practice 8:e70329.

8. Csárdi G, Nepusz T. 2006. The igraph software package for complex network research.

9. Danecek P, et al. 2021. Twelve years of SAMtools and BCFtools. GigaScience 10.

10. Flather CH, Hayward GD, Beissinger SR, Stephens PA. 2011. Minimum viable populations: is there a ‘magic number’ for conservation practitioners? Trends in Ecology & Evolution 26:307–316.

11. Formenti G, et al. 2022. The era of reference genomes in conservation genomics. Trends in Ecology & Evolution 37:197–202.

12. Garner BA, et al. 2016. Genomics in Conservation: Case Studies and Bridging the Gap between Data and Application. Trends in Ecology & Evolution 31:81–83.

13. Gobster P, Schneider, I., Floress, K., Haines, A. Arnberger, A. Dockry, M., Benton, Claire. 2021. Understanding the key characteristics and challenges of pine barrens restoration: insights from a Delphi survey of forest land managers and researchers. Restoration Ecology 29:e13273.

14. Goldstein J. 1978. THE PINE BARRENS: A STUDY OF ENVIRONMENTAL DECISION MAKING. Page 201. City University of New York, United States -- New York.

15. Goudet J. 2005. hierfstat, a package for r to compute and test hierarchical F-statistics. Molecular Ecology Notes 5:184–186.

16. Haines A, Farnsworth E, Morrison G 2011. New England Wild Flower Society’s Flora Novae Angliae A Manual for the Identification of Native and Naturalized Higher Vascular Plants of New England. Yale University Press.

17. Haller BC, Ralph PL, Messer PW. 2026. SLiM 5: Eco-evolutionary Simulations Across Multiple Chromosomes and Full Genomes. Molecular Biology and Evolution 43.

18. Jari O, et al. 2001. vegan: Community Ecology Package. The R Foundation.

19. Jombart T. 2008. adegenet: a R package for the multivariate analysis of genetic markers. Bioinformatics 24:1403–1405.

20. Kamvar ZN, Tabima JF, Grünwald NJ. 2014. Poppr: an R package for genetic analysis of populations with clonal, partially clonal, and/or sexual reproduction. PeerJ 2:e281.

21. Klütsch CFC, Laikre L. 2021. Closing the Conservation Genetics Gap: Integrating Genetic Knowledge in Conservation Management to Ensure Evolutionary Potential. Pages 51–82 in Ferreira CC, and Klütsch CFC, editors. Closing the Knowledge-Implementation Gap in Conservation Science: Interdisciplinary Evidence Transfer Across Sectors and Spatiotemporal Scales. Springer International Publishing, Cham.

22. Lacy, R.C., Miller, P.S., and Traylor-Holzer, K. 2025. Vortex 10 User’s Manual. 19 March 2025 update. IUCN SSC Conservation Planning Specialist Group, and Chicago Zoological Society, Apple Valley, Minnesota, USA.

23. Lacy, R.C., and Pollak, J.P. 2025. Vortex: A Stochastic Simulation of the Extinction Process. Version 10.10.0. Chicago Zoological Society, Brookfield, Illinois, USA.

24. Lee C, Robinson, G., Robinson, I., Lee, H. 2019. Regeneration of pitch pine (*Pinus rigida*) stands inhibited by fire suppression in Albany Pine Bush Preserve, New York. Journal of Forestry Research 30:233–242.

25. Lenth, R. 2024. emmeans: Estimated Marginal Means, aka Least-Squares Means. R package version 1.10.4.

26. Li H, Durbin R. 2009. Fast and accurate short read alignment with Burrows–Wheeler transform. Bioinformatics 25:1754–1760.

27. Likert R. 1932. A technique for the measurement of attitudes. Archives of Psychology 22 140:55–55.

28. Linton J. 2017. Recovery Strategy for the Karner Blue (Lycaeides melissa samuelis), Frosted Elfin (Callophrys irus) and Eastern Persius Duskywing (Erynnis persius persius) in Canada. Environment and Climate Change Canada.

29. Lüdecke D, Ben-Shachar M, Patil I, Waggoner P, Makowski D. 2021. “performance: An R Package for Assessment, Comparison and Testing of Statistical Models.” Journal of Open Source Software, 6(60), 3139. doi:10.21105/joss.03139.

30. Maschinski J, Albrecht MA. 2017. Center for Plant Conservation’s Best Practice Guidelines for the reintroduction of rare plants. Plant Diversity 39:390–395.

31. Mazerolle, M. J. 2023. AICcmodavg: Model selection and multimodel inference based on (Q)AIC(c). R package version 2.3.3.

32. Meyer, R. 2006. Lupinus perennis. In: Fire Effects Information System, [Online]. U.S. Department of Agriculture, Forest Service, Rocky Mountain Research Station, Fire Sciences Laboratory (Producer).

33. Michaels H, Cartwright, C., Wakeley Tomlinson, E. 2019. Relationships Among Population Size, Environmental Factors, and Reproduction in *Lupinus perennis* (Fabaceae). The American Midland Naturalist 182:160–180, 121.

34. NatureServe. 2026. NatureServe Explorer [web application].

35. New Hampshire Fish and Game Department. 2025. New Hampshire State Wildlife Action Plan. Concord, New Hampshire. https://www.wildlife.nh.gov/wildlife-and-habitat/nh-state-wildlife-action-plan/2025-state-wildlife-action-plan

36. Nielsen ES, Beger M, Henriques R, von der Heyden S. 2020. A comparison of genetic and genomic approaches to represent evolutionary potential in conservation planning. Biological Conservation 251:108770.

37. Ouborg NJ, Pertoldi C, Loeschcke V, Bijlsma R, Hedrick PW. 2010. Conservation genetics in transition to conservation genomics. Trends in Genetics 26:177–187.

38. Paris JR, Stevens JR, Catchen JM. 2017. Lost in parameter space: a road map for stacks. Methods in Ecology and Evolution 8:1360–1373.

39. Pavlovic N, Grundel, R. 2009. Reintroduction of Wild Lupine (*Lupinus perennis L*.) Depends on Variation in Canopy, Vegetation, and Litter Cover. Restoration Ecology 17:807–817.

40. Pedersen T. 2025. ggraph: An Implementation of Grammar of Graphics for Graphs and Networks. R package version 2.2.2, https://ggraph.data-imaginist.com.

41. Petitta IR, López-Uribe MM, Sabo AE. 2024. Biology and management of wild lupine (*Lupinus perennis L*.): a case study for conserving rare plants in edge habitat. Plant Ecology 225:373–389.

42. Petitta IR, Sabo AE, López-Uribe MM. 2025. Assessing the status of sundial lupine (*Lupinus perennis L.*)genetic diversity and population structure throughout its distribution. AoB PLANTS 17.

43. Plenzler M, Michaels, H. 2015. Seedling Recruitment and Establishment of *Lupinus perennis* in a Mixed-Management Landscape. Natural Areas Journal 35:224–234, 211.

44. Poland JA, Rife TW. 2012. Genotyping-by-Sequencing for Plant Breeding and Genetics. The Plant Genome 5.

45. R Core Team. 2024. R: A Language and Environment for Statistical Computing. R Foundation for Statistical Computing, Vienna, Austria.

46. R. Taylor H, Dussex N, van Heezik Y. 2017. Bridging the conservation genetics gap by identifying barriers to implementation for conservation practitioners. Global Ecology and Conservation 10:231–242.

47. Ragan L. 2019. Karner Blue Butterfly 5-Year Review.

48. Reed DH, O’Grady JJ, Ballou JD, Frankham R. 2003. The frequency and severity of catastrophic die-offs in vertebrates. Animal Conservation 6:109–114.

49. Reed MS, Graves A, Dandy N, Posthumus H, Hubacek K, Morris J, Prell C, Quinn CH, Stringer LC. 2009. Who’s in and why? A typology of stakeholder analysis methods for natural resource management. J Environ Manage 90:1933–1949.

50. Rivera-Colón AG, Catchen J. 2022. Population Genomics Analysis with RAD, Reprised: Stacks 2. Pages 99–149 in Verde C, and Giordano D, editors. Marine Genomics: Methods and Protocols. Springer US, New York, NY.

51. Robinson, D., Hayes, A., Couch, S., Hvitfeldt, E. 2025. broom: Convert Statistical Objects into Tidy Tibbles. R package version 1.0.11, https://broom.tidymodels.org/.

52. Sandström A, Lundmark C, Andersson K, Johannesson K, Laikre L. 2019. Understanding and bridging the conservation-genetics gap in marine conservation. Conservation Biology 33:725–728.

53. Schmidt C, Hoban S, Jetz W. 2023. Conservation macrogenetics: harnessing genetic data to meet conservation commitments. Trends in Genetics 39:816–829.

54. Shafer ABA, et al. 2015. Genomics and the challenging translation into conservation practice. Trends in Ecology & Evolution 30:78–87.

55. Shi XJ. 2004. Inbreeding and inbreeding depression in Lupinus perennis. PhD dissertation, Bowling Green State University, Bowling Green, Ohio.

56. Smallidge P, Leopold, D., Allen, C. 1996. Community Characteristics and Vegetation Management of Karner Blue Butterfly (*Lycaeides melissa samuelis*) Habitats on Rights-of-Way in East-Central New York, USA. Journal of Applied Ecology 33:1405–1419.

57. So, C., Hargreaves, A., Schoen, D. 2024. [Unpublished Lupinus perennis reference genome].

58. St.Clair JB, Richardson BA, Stevenson-Molnar N, Howe GT, Bower AD, Erickson VJ, Ward B, Bachelet D, Kilkenny FF, Wang T. 2022. Seedlot Selection Tool and Climate-Smart Restoration Tool: Web-based tools for sourcing seed adapted to future climates. Ecosphere 13:e4089.

59. USFWS. 2018. Species status assessment report for the frosted elfin (*Callophrys irus*), Version 1.2. Cortland, NY.

60. Weir BS, Cockerham CC. 1984. Estimating F-Statistics for the Analysis of Population Structure. Evolution 38:1358–1370.

61. Wickham H, Averick M, Bryan J, Chang W, McGowan LD, François R, Grolemund G, Hayes A, Henry L, Hester J, Kuhn M, Pedersen TL, Miller E, Bache SM, Müller K, Ooms J, Robinson D, Seidel DP, Spinu V, Takahashi K, Vaughan D, Wilke C, Woo K, Yutani H. 2019. “Welcome to the tidyverse.” Journal of Open Source Software, 4(43), 1686. doi:10.21105/joss.01686.

62. Wickham, H. 2023. modelr: Modelling Functions that Work with the Pipe. R package version 0.1.11, https://github.com/tidyverse/modelr, https://modelr.tidyverse.org.

