## Supplementary Materials for "LEAPING A CONSERVATION GENETICS GAP: COMBINING POPULATION GENETICS AND SOCIAL NETWORK ANALYSIS FOR *LUPINUS PERENNIS* RESTORATION"

APPENDIX I

PINE BARREN CONSERVATION MANAGEMENT SURVEY

**Start of Block: Consent Form**

Q1 This research project is being conducted by [REDACTED from the [REDACTED]. The purpose of this survey is to gain an understanding of your thoughts, opinions, and experience with genetic data in the scope of your professional career. We are hoping that you will participate in the study by completing the survey linked below. The questions should only take about 10-15 minutes of your time to complete and will benefit pine barren conservation. Your responses are voluntary and will be kept confidential. We do not collect contact information and all results will be generalized to ensure specific respondents cannot be identified based on demographics. If you have any questions about this survey, or if you have difficulties answering questions online, we will be happy to help and can be reached at the phone number or email addresses listed below. If you would prefer a paper copy of the survey, contact us and we will be happy to accommodate you. Questions about this project may be directed to: [REDACTED]

Q2 Do you consent to participate in this research survey?

- No (1)
- Yes (2)

*Skip To: End of Survey If Do you consent to participate in this research survey? = No*

**End of Block: Consent Form**

**Start of Block: Demographics**

Q3 Demographics

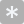

Q4
What organization do you work for?

________________________________________________________________

Q5 What is your job title?

________________________________________________________________

Q6 Do you work in the Northeast of the United States?

- No (1)
- Yes (2)

Q7 What is the highest degree you have received?

- High school diploma (1)
- Bachelor's degree (2)
- Master's degree (3)
- PhD (4)

Q8 In what year did you obtain your most recent degree?

________________________________________________________________

________________________________________________________________

________________________________________________________________

________________________________________________________________

________________________________________________________________

**End of Block: Demographics**

**Start of Block: Restoration Decisions**

Q9
The following questions ask you to consider restoration projects. A restoration involves assisting in the recovery of ecosystems that have been degraded or destroyed, as well as conserving the ecosystems that are still intact. 
Please consider any projects that fall under this definition, including population and species reintroduction and augmentation, and traditional ecosystem-level restorations.

Q10 Are you directly responsible for managing a population, species, or ecosystem?

- No (3)
- Yes (4)

Q11 If you were put in charge of a new restoration project, where would you consider sourcing seeds of an important plant species from?

- The area being restored (1)
- A connected nearby area (4)
- A disconnected nearby area (5)
- A far away area (6)
- I do not know how I would make this decision (7)
- Prefer not to answer (8)

Q12 Rank the factors that you consider most important for choosing a seed source:

______ Proximity of the source and restoration areas (1)

______ Similarity of climate between the source and restoration areas (2)

______ Size of the source population (4)

______ Genetic similarity between the source and restored populations (5)

______ Other (9)

**End of Block: Restoration Decisions**

**Start of Block: Implementation of Conservation Genetics**

Q13 Use of Genetic Resources The following questions will ask you about your opinions on and experiences with genetic resources. Please consider any data or analyses derived from DNA, RNA, or epigenetic sources when you answer the questions. Please consider research or recommendations you have used even if you did not work with or analyze the data yourself.

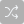

Q14 Rate your familiarity with the following DNA sequencing techniques

|  | Not at all familiar (1) | Somewhat familiar (2) | Moderately familiar (3) | Very familiar (4) | Extremely familiar (5) |
| --- | --- | --- | --- | --- | --- |
| Microsatellites (1) |  |  |  |  |  |
| Reduced representation (GBS, RAD, ddRAD, or DART-seq) (2) |  |  |  |  |  |
| Whole genome shotgun (6) |  |  |  |  |  |
| Low coverage whole genome shotgun (7) |  |  |  |  |  |
| Long read (Nanopore or PacBio) (8) |  |  |  |  |  |

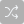

Q15 How useful do you think genetic data is for addressing:

|  | Not at all useful (65) | Slightly useful (66) | Somewhat useful (67) | Very useful (68) | Extremely useful (69) |
| --- | --- | --- | --- | --- | --- |
| Population connectivity (1) |  |  |  |  |  |
| Translocations (2) |  |  |  |  |  |
| Inbreeding (3) |  |  |  |  |  |
| Population or species restoration (4) |  |  |  |  |  |
| Captive breeding (5) |  |  |  |  |  |
| Taxonomic uncertainties (6) |  |  |  |  |  |

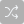

Q16 Check issues you have used genetic data to address:

- Population connectivity (9)
- Translocations (10)
- Inbreeding (11)
- Population or species restoration (12)
- Captive breeding (13)
- Taxonomic uncertainties (14)
- None of the above (15)

*Display this question:*

*If Check issues you have used genetic data to address: Is Less Than 6*

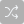

Q17 For issues you have not used genetic data for: Why not?

- Not appropriate (1)
- Insufficient funds (2)
- Did not have contact/staff who could do the work (3)
- Regulatory barriers (6)
- I did not know it was an option (5)
- I have not encountered these issues (8)
- Other (4) __________________________________________________

Q18 How likely would you be to use genetic resources if they were available for a future project?

- Extremely unlikely (1)
- Somewhat unlikely (2)
- Neither likely nor unlikely (3)
- Somewhat likely (4)
- Extremely likely (5)

Q19 Do you feel that you personally have enough information and knowledge to use genetic-based recommendations?

- Definitely not (1)
- Probably not (2)
- Might or might not (3)
- Probably yes (4)
- Definitely yes (5)

Q20 How often do you take genetics into consideration when writing or assessing species/ecosystem management plans?

- Never (1)
- Sometimes (2)
- About half the time (3)
- Most of the time (4)
- Always (5)
- I do not work with management plans (6)

Q21 Do you currently have anyone with genetic research training and/or experience on your team?

- No (4)
- Maybe (5)
- Yes (6)

Q22 How important is genetic diversity to population persistence?

- Not at all important (1)
- Slightly important (2)
- Moderately important (3)
- Very important (4)
- Extremely important (5)

**End of Block: Implementation of Conservation Genetics**

**Start of Block: Decision Making Networks**

*Display this question:*

*If Do you work in the Northeast of the United States? = Yes*

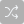

Q23 How often do you communicate with the following organizations when making either formal or informal decisions about plant conservation?

|  | Never (1) | Once per year (2) | 3-5 times per year (3) | Once per month (4) | Weekly (9) |
| --- | --- | --- | --- | --- | --- |
| US Environmental Protection Agency (60) |  |  |  |  |  |
| US Forest Service (1) |  |  |  |  |  |
| US Fish and Wildlife Service (20) |  |  |  |  |  |
| US Geological Survey (21) |  |  |  |  |  |
| Native Plant Trust (39) |  |  |  |  |  |
| Nature Conservancy (40) |  |  |  |  |  |
| Open Space Institute (58) |  |  |  |  |  |
| Maine Department of Agriculture, Conservation, and Forestry (59) |  |  |  |  |  |
| Massachusetts Department of Conservation and Recreation (49) |  |  |  |  |  |
| Massachusetts Division of Fisheries and Wildlife (50) |  |  |  |  |  |
| Nantucket Conservation Foundation (52) |  |  |  |  |  |
| Nantucket Land Bank (53) |  |  |  |  |  |
| Southeast Massachusetts Pine Barren Alliance (54) |  |  |  |  |  |
| New Hampshire Fish and Game (44) |  |  |  |  |  |
| New Hampshire Natural Heritage Bureau (45) |  |  |  |  |  |
| Pinelands Preservation Alliance (55) |  |  |  |  |  |
| Albany Pine Bush (46) |  |  |  |  |  |
| New York Department of Environmental Conservation (47) |  |  |  |  |  |
| New York Department of Parks, Recreation, and Historical Preservation (48) |  |  |  |  |  |
| Wilton Wildlife Preserve (51) |  |  |  |  |  |
| Vermont Agency of Natural Resources (56) |  |  |  |  |  |
| Vermont Fish and Wildlife Department (57) |  |  |  |  |  |

Q24 Do you communicate formally or informally with organizations other than those listed above when making plant conservation decisions?

- No (1)
- Yes (2)

**End of Block: Decision Making Networks**

**Start of Block: Decision Making Networks Open Response**

*Display this question:*

*If Do you communicate formally or informally with organizations other than those listed above when m... = Yes*

Q25 List and rate how often you communicate with organizations not listed in the last question when making formal or informal decisions about plant conservation.

|  | Never (1) | Once per year (2) | 3-6 times per year (3) | Once per month (4) | Weekly (5) |
| --- | --- | --- | --- | --- | --- |
| Organization 1 (6) |  |  |  |  |  |
| Organization 2 (7) |  |  |  |  |  |
| Organization 3 (8) |  |  |  |  |  |
| Organization 4 (9) |  |  |  |  |  |
| Organization 5 (10) |  |  |  |  |  |
| Organization 6 (11) |  |  |  |  |  |
| Organization 7 (12) |  |  |  |  |  |
| Organization 8 (13) |  |  |  |  |  |
| Organization 9 (14) |  |  |  |  |  |
| Organization 10 (15) |  |  |  |  |  |

**End of Block: Decision Making Networks Open Response**

APPENDIX II

GENOTYPE-BY-SEQUENCING PROTOCOL

**Double Digest MluC1 Msp1 RAD-seq**

**Digestion -> Ligation -> Clean, concentrate, and size selection -> PCR -> Clean and concentrate -> quantify and pool -> size select with tech reps -> QAQC**

**I. Double Digestion**

1. Before starting, quantify the sample using the QuBit quantification and normalize all samples to 8.75 ng/ul at 36ul final volume in a 96-well plate.
2. Add 12 ul normalized DNA (8.75 ng/ul) of each sample to a new plate
3. Make the mastermixes for the enzyme pair in 1.5 ml tubes. Keep the enzymes cold (on ice or in a freezer block) at all times.
4. Aliquot 8 ul of mastermix into each well for a total final volume of 20 ul.
5. Place on the thermocycler and run at 37°C for 3 hours. Deactivate enzymes by incubating the reactions at 65°C for 20 minutes, and then hold for 4°C. It is a good idea to proceed to ligation quickly.

| **DOUBLE DIGESTION MASTER MIX** | **1 sample (µl)**  **for 12 µl DNA** | **100 samples (µl)** |
| --- | --- | --- |
| H_2_O, pH 7 **OR** 10mM Tris-HCl, pH 8 | 5 | 500 |
| NEB 10X Cutsmart buffer | 2.0 | 200 |
| NEB MluC1 | 10 units: 0.5 | 50 |
| NEB Msp I | 10 units: 0.5 | 50 |
| **Total** | **8.0** | **800** |

**II. Adapter Ligation**

The adapters are as follows:

| MspI_P1.1 | ACACTCTTTCCCTACACGACGCTCTTCCGATCT | 100 nmol custom oligo with HPLC purification |
| --- | --- | --- |
| MspI_P1.2 | /5Phos/CGAGATCGGAAGAGCGTCGTGTAGGGAAAGAGTGT | 100 nmol custom oligo with HPLC purification |
| MluCI_P2.1 | GTGACTGGAGTTCAGACGTGTGCTCTTCCGATCT | 100 nmol custom oligo with HPLC purification |
| MluCI_P2.2 | /5Phos/AATTAGATCGGAAGAGCGAGAACAA | 100 nmol custom oligo with HPLC purification |

The lowest scale for the oligos as HPLC purified duplexed DNA will be 100 nmol which will give a minimum yield of 3.8 nmol. We don’t know the exact concentration and volume (since the document mentions 4uM as dummy value) but if going by 4 uM concentration and 2 ul per reaction, this should be enough for roughly 475 reactions. The adapters are already annealed and purified (from IDT).

MluC1 is the frequent cutter and Msp1 is the infrequent cutter

If we are skipping speedbead cleaning after digestion, perform the ligation immediately after digestions and use the same buffer as used in the double digestion and add ATP (the buffer that comes with the ligase already has ATP, but we have to add it if using CutSmart). Use 1ul of P1 and P2 at 10uM (or 0.4ul of 4uM). If we want to follow Peterson’s protocol for the amount of adapter per cutting site, go to Mayra’s copy of Brent’s calculation sheet <https://docs.google.com/spreadsheets/d/1aYz11M1c8EnA_IeH1_rd1AuJZuRoWP-ZPTC5L3dSzCY/edit#gid=0>

Following the above calculation, we would need 0.4ul for the frequent cutter (P2) and 2ul for the infrequent cutter (P1). It is good to bias ligation toward the infrequent cutter to reduce frequent-frequent fragments.

1. Dilute the adaptors to 4um in new tubes (redo this calc each time you get new adaptors from IDT):
   a. P1: 25ul adaptor (10um) into 600ul water
   b. P2: 5.8ul adaptor (10um) into 139.2ul water
2. Spin the plates after digestion using the centrifuge
3. Make the ligation master mix as below in two 1.5ml tubes. Keep the T4 DNA ligase, ATP, and mastermix on ice at all times. Add the T4 DNA ligase last, right before aliquoting it into the wells of the digested DNA plate
4. Add 20 ul of mastermix to each well using a repeater pipette. Spin down the plate, then mix each well using a multichannel

| **LIGATION MASTER MIX** | **Catalogue Number** | **1 sample**  **(µl)** | **100 samples (µl)** |
| --- | --- | --- | --- |
| H_2_O, pH 7 **OR** 10mM Tris-HCl, pH 8 |  | 11.1 | 1,110 |
| NEB 10X Cutsmart buffer | B7204 | 2.0 | 200 |
| 10 mM ATP | P0756 | 4.0 | 400 |
| P1 adaptor (4 uM) |  | 2 | 200 |
| P2 adaptor (4 uM) |  | 0.4 | 40 |
| NEB T4 DNA Ligase | M0202 | ~200 U: 0.5 | 50 |
| **Total** |  | **20** | **2000** |

Incubate at 22°C for more than 3 hours. I (Cooper) do this overnight for 999 minutes (max on the thermocycler), then place at 65°C for 10-20 minutes, and hold at 4°C

Spin the plate down after incubation and move to clean and concentrate immediately.

**III. Clean and Concentration**

After ligation, purify and concentrate the individual reaction in a 96-well plate, using SPRI beads. To each well add 64 µl (1.6x) of beads using a multichannel pipette (aliquot the beads beforehand in a 8-strip of PCR tubes or a reservoir, and then dispense them from there) and then proceed using the 96-well magnet. All steps use multichannel pipettes:

- Spin ligated plate down to collect condensation before removing foil
- Let beads come to room temperature and mix well before pipetting
- Add 64 µl of SPRI beads to each well, mix, let stand for 5 mins
- Move the plate to the magnet, let stand for 2-5 mins
- Discard supernatant (beads hold the DNA against the magnet)
- Keeping plate on the magnet, add 200 µl of 80% EtOH to each well
- Wait 15-30 seconds, then remove the ethanol
- Repeat for a second 80% EtOH wash
- Add 200ul 100% EtOH to each well
- Wait 15-30 seconds and remove all the ethanol
- Let the plate dry completely for 5-10 mins
- Remove plate from magnet and resuspend each well in 12 µl of 10 mM Tris-HCl pH 8.0
- Mix resuspended wells by pipetting and let stand for 2 mins
- Put the plate back on the magnet, let stand for 2 mins
- Transfer 10 µl of the supernatant (containing your DNA) to a clean plate.

**IV. Amplification**

Barcodes are added at this stage using IDTs XGen UDI Primer Plates (Plate 1, 8nt: product# 10005922; Plate 2, 8nt: product# 10009816). Each well of the Primer Plate contains 20 mM of barcode primers (10 mM of each i5 and i7 primers). Aliquot 1 µl of barcoded primers (set 1–96 or 97–192 at 0.4 ng/µl) into the double digestion plate. DO NOT transfer the volume of digested samples directly into the plate of barcoded primers, IDT provides much more than 1 ul. Using only 1 ul of primers also allows you to split a plate across multiple reactions with relative success and also allows you to retry an amplification by not using all of your ligated sample. Regardless of your strategy, the key is for each sample to contain the same concentration of primers so that amplification occurs evenly across samples.

I (Cooper) recommend aliquoting the DNA template by multichannel, then add HotStart by repeater, then spin the plate down before adding the primers and mixing the reaction solution together with a repeater. Saves tips and time. Make sure to keep the plate orientations the same before and after transfer. One thing to keep in mind is that increasing the number of PCR cycles increases the likelihood of PCR bias significantly affecting your sequencing data.

| **AMPLIFICATION MIX** | **1 rxn (µl)** | **1 library**  **(100 rxns)** |
| --- | --- | --- |
| DNA template (purified ligation) | 4 | Pulled from digestion plate |
| Kapa HIFI HotStart 2X | 5 | X100 = 500 |
| 20 mM Illumina Primer Pair | 1 | Pulled from primer plate |
| Total | 10.0 |  |

| Amplification cycles | ^0^C | time |
| --- | --- | --- |
| Step 1. | 98 | 00.00.30 |
| Step 2. | 98 | 00.00.30 |
| Step 3. | 62 | 00.00.20 |
| Step 4. | 72 | 00.00.30 |
| repeat steps 2-4 12–16X |  |  |
| Step 5. | 72 | 00.05.00 |
| Step 6. | 4 | forever |

**V. Clean and Pool**

At this point, the barcodes are attached to the sequences, if all has gone well. We need to clean and partially size select our samples to get rid of the polymerase and any latent primer dimers. To each well add 0.80x concentration of SPRI beads using a multichannel pipette (aliquot the beads beforehand in a 8-strip of PCR tubes or reservoir and then dispense them from there) and then proceed using the 96-well magnet. All steps use multichannel pipettes:

- Spin amplification plate down to collect condensation before removing foil
- Let beads come to room temperature and mix well before pipetting
- Add 8.0µl of SPRI beads to each well, mix, let stand for 5 mins
- Move the plate to the magnet, let stand for 2-5 mins
- Discard supernatant (beads hold the DNA against the magnet)
- Keeping plate on the magnet, add 200 µl of 80% EtOH to each well
- Wait 15-30 seconds, then remove the ethanol
- Repeat for a second 80% EtOH wash
- Add 200ul 100% EtOH to each well
- Wait 15-30 seconds and remove all the ethanol
- Let the plate dry completely for 5-10 mins
- Remove plate from magnet and resuspend each well in 15 µl of 10 mM Tris-HCl pH 8.0
- Mix resuspended wells by pipetting and let stand for 2 mins
- Put the plate back on the magnet, let stand for 2 mins
- Transfer 13 µl of the supernatant (containing your DNA) to a clean plate.

Qubit all of the samples that will be sequenced in the same library and renormalize to the lowest concentration. The idea is to represent each sequence equally in the final library, which may involve normalizing and pooling across plates. It is safe to stop once you have cleaned out the enzyme, so you can wait until all of your samples are done to renormalize.

**VI. Size Selection**

Using the SimRAD protocol, you can determine the number of likely fragments at a given size range. Depending on your intent for the end results, choose a specific range of sizes. We will go with ~300bp. The size can be selected either by the Pippin Prep (or other similar equipment), by running a gel and cutting it with a razor blade and extracting the appropriate range, or by using SpeedBeads (<https://www.zoology.ubc.ca/~rieseberg/RiesebergResources/ampurexp-beads-dna-clean-up-and-size-selection-dan-e/>).

The pooling after the PCR should have gotten rid of smaller than 300 bp, if we were using ~0.85X (according to Rieseberg’s site) beads. Now, we want to get rid of the larger fragments. To do that, we select a concentration of beads that will only allow fragments larger than ~700 bp bind. Note the elution volume at the end is up to you. I (Cooper) find that anything less than 1/10 of the volume of the beads makes it difficult to resuspend, so I would recommend using that as your floor rather than 12 ul.

- Split the pooled library into 3-5 technical replicate 2ml tubes
- Let beads come to room temperature and mix well before pipetting
- Add 0.60x beads to each replicate tube and pipette to mix
- Let stand 5 mins
- Place on plate magnet and allow pellet to form (~2 mins)
- Transfer supernatant (containing smaller fragments that did not bind to the beads at 0.60x) from each tube to a fresh tube off the magnet
- Mix beads well. Add 1x concentration of beads to each tube. Mix and let stand 5 mins
- Place tubes on magnet and allow pellet to form (~2 min)
- Remove and discard supernatant
- Wash with 80% EtOH, let sit 30 secs, then remove supernatant
- Repeat for 2 80% EtOH washes
- Wash with 100% EtOH, let sit 30 secs, remove as much EtOH as possible
- Let tubes dry for 5-10 mins with caps open
- Remove tubes from magnet & elute in 15 ul 10mM Tris. Mix well
- Let stand for 5 minutes off magnet
- Place on magnet and allow pellet to form (~2 min)
- Carefully transfer 12 ul supernatant to fresh tubes
- Recombine technical replicates

Qubit, nanodrop, and run the library on a gel for QAQC. If all looks good, send for sequencing.

APPENDIX III

*VORTEX* SIMULATION INPUT PARAMETERS

Populations simulated for 160 years for 250 iterations

Scenario Settings Notes: This simulates the founder effects of new populations of uniform size pairing NH populations as seed sources mixed with Albany, NY seed. The new population is founded in year 2 with equal numbers of plants from each NH population and the paired NY population. We examine genetic diversity over ~40 generations, which is the same time period used for MVP.

Population extinction is defined as having fewer than 5 individuals at the end of 160 years, and we have the simulation catalog gene diversity by year and iteration.

Sequence of events in each time cycle:

EV

Breed

Mortality

Age

Disperse

Harvest

Supplement

rCalc

Ktruncation

GSUpdate

PSUpdate

ISUpdate

Census

Extinction defined by: N<5

Species Description Notes: Currently we do not quantify inbreeding depression, although this may be possible with values from So et al.'s work on genetic load in the species.

No inbreeding depression.

State Variables Notes: No state variables.

Correlation of EV among populations = 0.5

Dispersal Notes: No dispersal between populations (translocations are coded in directly later).

Reproductive System:

Reproductive System Notes: The plant is hermaphroditic with the age of first offspring at 3 years, maximum lifespan and reproduction at 10 years, 1 brood per year, and a maximum number of offspring of 500 per brood.

Offspring are not dependent on their dam and there is currently no estimate of density dependence in the species, although this could potentially be loosely estimated using Michaels et al.'s work in Michigan and Ohio populations.

Hermaphroditic breeding

Females breed from age 3 to age 10

Males breed from age -1 to age 10

Maximum age of survival: 10

Sex ratio (percent males) at birth: 50

Correlation of EV between reproduction and survival = 0.5

EV sampled from Beta distributions.

Reproductive Rates Notes: Mean and SD number of offspring is estimated from lupine management survey and Shi et al.

Mortality Notes: Mortality rate parameters were calculated in Shi et al for the first year and Zaremba et al for following years. Both values are reported in the 2003 Karner Blue Butterfly Recovery Plan.

Catastrophe Notes: Frankham et al. estimates that organisms experience one catastrophe that results in 50% die off per generation, which calculates to a 5% chance per year given lupine's lifespan. Reproductive penalties for such a rare drought appear in the USDA page for L. perennis

Catastrophes are made global so simulations across populations are comparable.

Mate Monopolization Notes: Michaels estimates a 10% selfing rate in natural populations

Initial Population Size Notes: Population sizes are set as first year individuals for newly founded population.

Single populations are not supplemented and should be started at MVP.

Carrying Capacity Notes: Carrying capacity set to the known MVP of the population

Harvest Notes: Harvest and translocation pages set to move 12 plants from the seed bank to the wild population ever two years until the wild population reaches the minimum viable population size. Assuming 100% survival of translocations because these are transplants grown in the greenhouse and cared for after transplant, which should result in low mortality and potentially additional supplementation in off years to make up for lost individuals.

Supplementation Notes: Supplementing an equal number of individuals from the AL population to bring the founders up to MVP size.

Single populations are not supplemented.

Percent of adult females breeding each year: 65.7

with EV(SD): 16.9

Percent selfing: 10

Normal distribution of brood size with mean: 128 with SD: 155

Female annual mortality rates (as percents):

Age 0 to 1: 85 with EV(SD): 5

Age 1 to 2: 50 with EV(SD): 10

Age 2 to 3: 50 with EV(SD): 10

After age 3: 50 with EV(SD): 10

Catastrophe 1: Catastrophe1

Global impact

Frequency (%): 5

Reproduction reduced by severity multiplier: 0.66

Survival reduced by severity multiplier: 0.5

Initial population size:

Age	0	1	2	3	4	5	6	7	8	9	10	Total

Females	0	36	0	0	0	0	0	0	0	0	0	36

Carrying capacity: 71

with EV(SD): 0

Supplementation from year 1 through year 1 by increments of 1

Age	0	1	2	3

Females	0	0	35	0

Genetics options:

Genetics Notes: Allele frequencies estimated from GBS data of the wild populations. Allele frequencies are calculated for a total of 12,433 biallelic loci.

Mutation applies to all loci equally as they should all be neutral. Mutation rate for Fabaceae is estimated at 4.73E-9/year. Assuming a generation time of 4 years, the probability of a mutation occuring in a single neutral locus is 1.90E-8/generation.

Supplements are drawn from the last population, order of populations in gene file must match order of listed populations.

12433 nuclear loci (and one mtDNA locus) simulated

Genetic summary statistics based only on loci after locus 1

First locus is default with infinite alleles model

Mutation modeled for 12433 loci

with mutation rate = 1.9E-08
